# A systematic mapping of the genomic and proteomic variation associated with monogenic diabetes

**DOI:** 10.1101/2023.01.19.524722

**Authors:** Ksenia Kuznetsova, Jakub Vašíček, Dafni Skiadopoulou, Janne Molnes, Miriam Udler, Stefan Johansson, Pål Rasmus Njølstad, Alisa Manning, Marc Vaudel

## Abstract

**Aims:** Monogenic diabetes is characterized as a group of diseases caused by rare variants in single genes. Multiple genes have been described to be responsible for monogenic diabetes, but the information on the variants is not unified among different resources. In this work, we aimed to develop an automated pipeline that collects all the genetic variants associated with monogenic diabetes from different resources, unify the data and translate the genetic sequences to the proteins.

**Methods:** The pipeline developed in this work is written in Python with the use of Jupyter notebook. It consists of 6 modules that can be implemented separately. The translation step is performed using the ProVar tool also written in Python. All the code along with the intermediate and final results is available for public access and reuse.

**Results:** The resulting database had 2701 genomic variants in total and was divided into two levels: the variants reported to have an association with monogenic diabetes and the variants that have evidence of pathogenicity. Of them, 2565 variants were found in the ClinVar database and the rest 136 were found in the literature showing that the overlap between resources is not absolute.

**Conclusions:** We have developed an automated pipeline for collecting and harmonizing data on genetic variants associated with monogenic diabetes. Furthermore, we have translated variant genetic sequences into protein sequences accounting for all protein isoforms and their variants. This allows researchers to consolidate information on variant genes and proteins associated with monogenic diabetes and facilitates their study using proteomics or structural biology. Our open and flexible implementation using Jupyter notebooks enables tailoring and modifying the pipeline and its application to other rare diseases.

**Research in context:**

- Monogenic diabetes is a group of Mendelian diseases with an autosomal-dominant pattern of inheritance.
- Monogenic diabetes is mainly caused by rare genetic variants that are usually evaluated manually.
- The data on the variants are stored in several resources and are not unified in terms of the genomic coordinates, alleles, and variant annotation.
- What can be done for the systematic evaluation of the variants and their protein consequences?
- In this work, we have created an automated Jupyter notebook-based pipeline for the collection and unification of the variants associated with monogenic diabetes.
- The database of the genetic variants was created and translated to all possible variant protein sequences.
- These results will be used for the analysis of proteomics data and protein structure modeling.

## Introduction

The most common forms of monogenic diabetes are maturity-onset diabetes of the young (MODY), neonatal diabetes (1), inherited lipodystrophies, mitochondrial diabetes, among others (2). Today, international guidelines are available for the diagnostic and follow-up of patients with suspected MODY (3). These patients may now receive a molecular genetic diagnosis using diagnostic gene sequencing panels (panelapp.genomicsengland.co.uk/panels/472). This allows precise MODY subtyping and, depending on the diagnosis, the opportunity to avoid lifelong insulin medication and complications through lifestyle management or alternative treatment using oral antidiabetic drugs (4). Furthermore, because the early correct diagnosis may implement successful treatment with low doses of sulfonylurea or diet alone and postpone complications, the timeline of diagnosis and care is thus crucial (4). It is estimated that around 77 % of monogenic diabetes cases remain undiagnosed (5). These patients and their relatives remain unaware of their familial condition and do not benefit from adapted care.

The first challenge in establishing a firm diagnosis in all cases of MODY is the mapping of all genes that can cause monogenic diabetes. To date, multiple genes have been discovered to have associations with familial forms of diabetes (2). Fourteen of them are notably often referred to as “the MODY genes” (3), although this list is subject to debate in the literature (6) and is being systematically assessed by international experts using established guidelines to determine gene-disease relationships (clinicalgenome.org/affiliation/40016). A second challenge is the difficulty to evaluate the pathogenicity of genetic variants (7), for which there is also an ongoing international effort to establish guidelines and provide expert variant curations in the ClinVar database (clinicalgenome.org/affiliation/50016) (8). Furthermore, the response to alternative treatment might differ between populations (9). One of the ways to address the challenge of precise diagnostics is to complement genetic screening with additional data, combining both molecular and clinical dimensions (10).

The recent advent of high-performance computational models for protein structures notably holds the promise to democratize the study of gene sequences (11). We aim to study the properties of the proteins encoded by genes carrying alleles suspected to cause monogenic diabetes. This can, for example, help us understand whether the structure and properties of the protein are affected by these alleles, hence shedding light on the pathogenicity of the variant investigated (12).

The adoption of these approaches is impaired by the difficulty of mapping variants associated with monogenic diabetes to the different forms of proteins that they encode. First, given that the variants associated with monogenic diabetes are rare, the coverage by genomic databases is low. Maintaining an updated list of variants requires constant monitoring of the literature by experts. Second, variants reported in the literature often lack standardization in their identifiers and coordinates, making it challenging to map them to a given genome build and requiring manual variant mapping. Third, inferring the consequences on protein sequences is still a daunting task for some variants (those alleles affecting splice sites and untranslated regions [UTRs], for example). Fourth, a given protein-coding sequence might encode different protein isoforms, which will produce different forms of proteins upon folding and post-translational modification (13), hence for a given variant multiple protein sequences need to be investigated. Mapping genetic variants associated with monogenic diabetes from genes to proteins is therefore not tractable and sustainable without automation using dedicated bioinformatic tools.

Here, we describe a new open-source modular pipeline based on Jupyter notebooks (eprints.soton.ac.uk/403913) that allows for the systematic collection of variants associated with monogenic diabetes and their mapping to Ensembl (14) and ClinVar (8). We demonstrate how the different genes associated with monogenic diabetes harbor variants of different levels of pathogenicity. Finally, we port the variant sequences to the protein level and provide the resulting sequences in a standard format that can readily be used for proteomic and structural proteomic analyses using mass spectrometry or protein structure modeling.

## Methods

### General architecture

The pipeline consists of seven independent modules written in Python using Jupyter notebooks (figure 1). The notebooks are chained together as a pipeline, but they can also be used as standalone applications, or integrated into other pipelines. First, the pipeline takes a list of genes and extracts exonic variants from Ensembl retaining only those variants that are predicted to affect protein sequences. Next, the program integrates the variants from ClinVar and maps them to Ensembl. Similarly, variants are extracted from the literature, here using the literature mining by Rafique et al. (15), and mapped to Ensembl. Subsequently, the harmonized collection of variants is consolidated in a database stored in the form of a text file that can easily be parsed and reused. Finally, the table of variants is mapped to all the transcripts linked by Ensembl to the genes of interest to obtain protein sequences of all the possible isoforms encoded by these genes. In this last step, the DNA sequences are translated to amino acid sequences and stored as protein FASTA files. To visually inspect the results, a separate module allows overlaying all the variants that can possibly affect a given gene onto the corresponding amino acid sequences.

**Figure 1.**
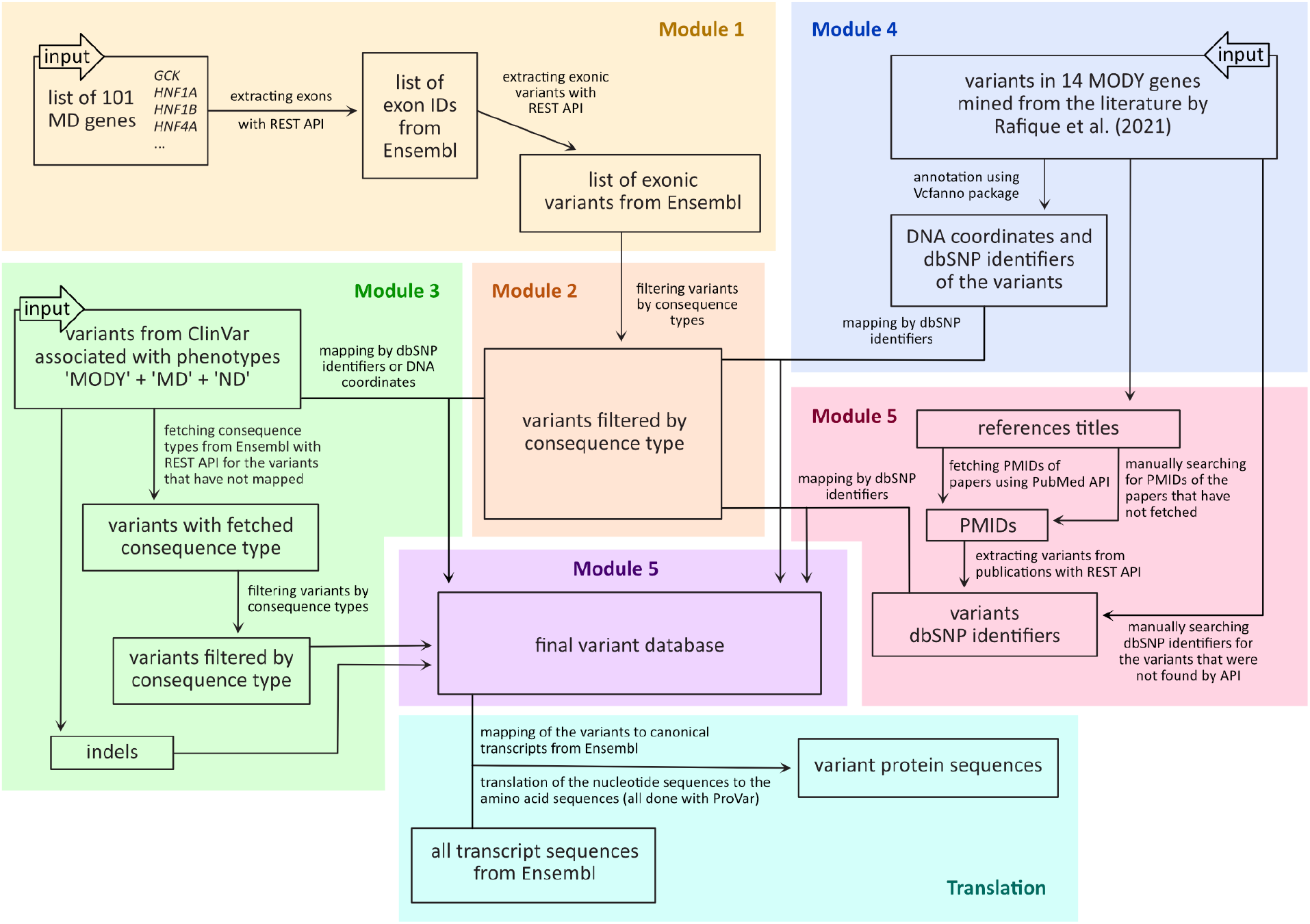
General architecture of the pipeline. *Module 1*: extract variants from Ensembl affecting genes associated with monogenic diabetes. *Module 2*: filter variant by consequence on the protein. *Module 3*: extract variants from ClinVar affecting genes associated with monogenic diabetes. *Modules 4* and *Module 5*: extract the variants from the literature using the mining by Rafique et al. *Module 6*: consolidate the variants in a single table. *Translation*: produce the possible variant protein sequences. MODY - maturity-onset diabetes of the young, MD - monogenic diabetes, ND - neonatal diabetes, API - application programming interface, PMID - identifiers of scientific publications from the PubMed database, dbSNP identifiers - identifiers of the genomic variants from the dbSNP database, that start from “rs”.

### Module 1 - Mining variants in Ensembl

In *Module 1*, a list of genes is mapped to Ensembl genes, transcripts, and exon identifiers using the Ensembl REST API (16). Subsequently, the Ensembl REST API is queried using the exon identifiers to return all variants in Ensembl overlapping with the corresponding regions along with their annotation (identifiers, coordinates, consequences, etc.). For multi-allelic variants, we treat every alternative allele as a different entry, and add it as a new line to the table produced. The example of the top rows of the reference table is given in the Supplementary materials, table 1.

### Module 2 - Categorizing by consequence and pathogenicity

*Module 2* takes as an input the table of exonic variants from Ensembl generated in *Module 1*. For each consequence type, the prevalence of variants with different levels of pathogenicity is computed and visualized as heat maps (figure 2A). The same is done for the data from ClinVar with all the variants regardless of the associated phenotype (figure 2B).

**figure 2.**
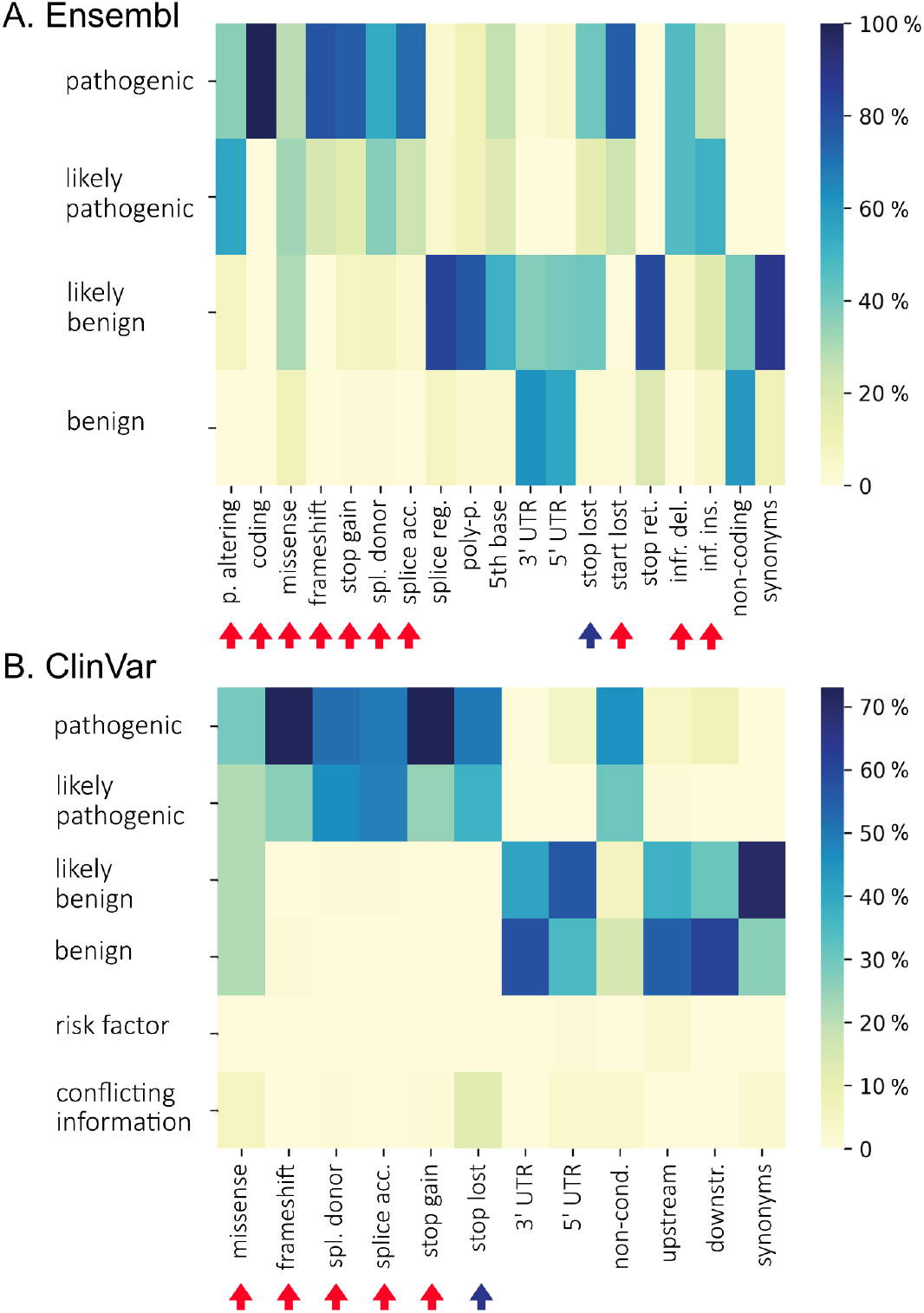
Distribution of the consequence types within pathogenicity categories of the variants from Ensembl (A) and ClinVar (B). The heatmap shows the percentage of variants within pathogenicity categories for each consequence type. Arrows show the consequence types that were left in the Ensembl reference table created in *Module 2* after filtering. “Stop lost” is marked with a blue arrow as it has conflicting evidence in two sources (see detailed in the text).

By default, only variants whose pathogenicity is classified mostly as *likely pathogenic* or *pathogenic* are retained for further analyses: ‘missense_variant’, ‘protein_altering_variant’, ‘coding_sequence_variant’, ‘frameshift_variant’, ‘splice_donor_variant’, ‘splice_acceptor_variant’, ‘splice_donor_5th_base_variant’, ‘start_lost’, ‘stop_gained’, ‘stop_lost’, ‘inframe_deletion’, ‘inframe_insertion’.

### Module 3. Mapping of the variants from ClinVar

*Module 3* takes as an input three tables exported from ClinVar containing all the variants associated with three phenotypes: “MODY”, “monogenic diabetes”, and “neonatal diabetes”. A few top rows of one of these tables are given in the Supplementary materials table 2 as an example. ClinVar annotates variants with different levels of pathogenicity: “pathogenic”, “likely pathogenic”, “uncertain significance”, “likely benign” or “benign”. The variants presenting dbSNP identifiers are passed to *Module 2*. The others are mapped to Ensembl using genomic coordinates and alleles and predicted consequences are obtained with the Ensembl Variant Effect Predictor (VEP)(17) called on all variants using the Ensembl REST API. The variants with predicted consequences according to Ensembl are filtered in the same way as in *Module 2*. The variants where no consequence type was returned were all either short deletions, short insertions, or other short fragment replacements. The table with the example of such variants is represented as table 3 in the Supplementary materials. This subset is directly passed to *Module 6*.

### Module 4. Mapping variants from the literature (step 1 of 2)

*Module 4* takes variants associated with monogenic diabetes according to the literature to cover the variants where the association with monogenic diabetes is not yet consolidated in ClinVar or Ensembl. As input we used the variants mined in the review by Rafique et al. (15) and provided as a supplementary table. The module uses the Vcfanno (18) library to annotate the variants with DNA coordinates and the dbSNP identifiers where possible. The Ensembl REST API is queried as in *Module 1*, and the results are passed to *Module 6*.

### Module 5. Mapping variants from the literature (step 2 of 2)

For the variants that could not be mapped automatically in *Module 4*, the title of the publication as obtained from *Module 4* is queried against the Entrez Programming Utilities API (www.ncbi.nlm.nih.gov/books/NBK25500) to return the PubMed (pubmed.ncbi.nlm.nih.gov) identifiers (PMIDs) of these articles. Note that some of the PMIDs were mapped and had to be added manually. Next, these PMIDs are used to query the same API and return the “rs” identifiers of the variants mentioned in these publications. Finally, the variants are mapped to Ensembl using their identifiers and passed to *Module 6*. Note that not all these variants could be mapped automatically and those that did not map to Ensembl were formatted manually for input to *Module 6*.

### Module 6 - Consolidation as table and VCF file

*Module 6* combines all the tables produced by the previous modules and creates Venn diagrams showing the number of variants obtained from the different sources and their overlap. The example of tables used as an input here is given as table 4 in Supplementary materials. This table is then used to create a Variant Calling Format (VCF) file listing the site of all the variants.

We further categorized the variants in two levels of pathogenicity evidence. *Level 1* variants include the variants from ClinVar obtained in *Module 3* associated with three phenotypes: “MODY”, “monogenic diabetes”, or “neonatal diabetes” regardless of their reported pathogenicity, complemented with the variants extracted from the literature in *Module 4* and *Module 5. Level 2* variants undergo stricter filtration criteria. For the variants obtained from ClinVar in *Module 3*, all variants labeled as *benign, likely benign*, or of *unknown pathogenicity* were filtered out. For the variants obtained from the literature in *Module 4* and *Module 5*, variants in *BLK, KLF11*, and *PAX4* were removed as these genes were reported to lack association with MODY in more recent literature (6).

### Translation of the variant sequences into protein sequences

The translation step was performed using the ProVar tool (github.com/ProGenNo/ProHap).

Shortly, the variants from the table obtained in *Module 6* were mapped to canonical cDNA sequences from Ensembl to retrieve all the transcripts of the same gene with the annotation of start and stop codons. After mapping, all the sequences were translated to their amino acid sequences and written into a protein FASTA file. Both the table output and FASTA examples are given in Supplementary materials as table 5 and FASTA 1.

### Sequence overlay

All the variants from the resulting database were overlaid with the reference protein sequences obtained from Ensembl for all the transcripts of all genes. The variants are represented using two separate rows corresponding to the two levels of pathogenicity confidence (see example in figure 3). FASTA files are parsed using the Pyteomics library (19). The protein sequences and variants are plotted using the Matplotlib library (20).

**figure 3.**
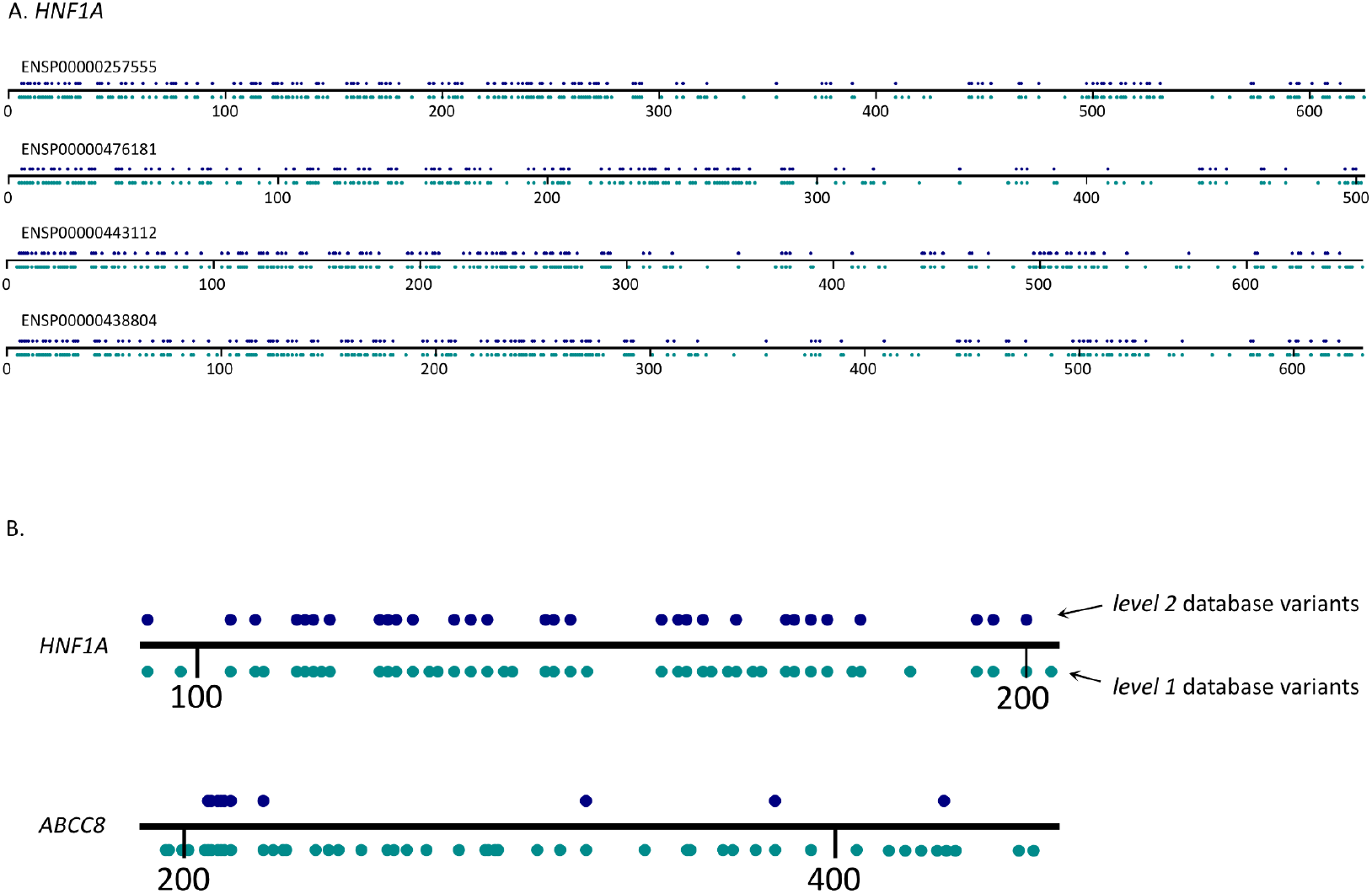
A: All isoforms of the protein product of *HNF1A* with the amino acid variants from the database mapped on the sequence. B: Examples of the random fragments of the products of the canonical isoforms of *HNF1A* and *ABCC8* showing the difference in density of variants from *level 1* and *level 2* databases. The top row of blue dots represents the positions of the variants from the *level 2* database, whilst the bottom green row represents the positions of the variants from the *level 1* database. Being the most well-studied gene, *HNF1A* has an almost equal number of dots in both rows as most of the reported variants are revised and confirmed as pathogenic. *ABCC8* has a lot of variants in the *level 1* database that are not confirmed as pathogenic and, thus, are not in the *level 2* database.

## Results

Unlike more common forms of diabetes like type 2 diabetes (T2D), where large numbers of samples are available and federated initiatives consolidate information on genetic variants and their consequences in aggregated and harmonized forms (e.g. t2d.hugeamp.org), monogenic diabetes, as a rarer disease, relies on small cohorts and information on genetic variants is scattered in the literature and online databases. The aggregation and comparison of variants known to be associated with monogenic diabetes therefore currently relies on expert manual curation and annotation.

### Mining variants in genes associated with monogenic diabetes

We mined variants based on a list of 101 genes associated with monogenic diabetes, aiming at being as comprehensive as possible. These 101 genes consist of: i. 14 MODY genes taken from OMIM (21) or other reviews on MODY such as (3); ii. 10 genes associated with neonatal diabetes, lipodystrophy, and insulin signaling taken from (2); iii. 77 genes having variants with any evidence of association with either MODY, neonatal diabetes, or just the condition referred to as “monogenic diabetes” by ClinVar.

### Categorizing by consequence and pathogenicity

Since monogenic diabetes, being a Mendelian disease, is determined mostly by rare, highly penetrant coding variants (5), we focused on exonic variants when mining Ensembl. Nevertheless, some variants reported in ClinVar and in the literature mapped to untranslated regions (UTR) and splice regions. In these cases we decided whether to keep these variants or not based on the pathogenicity of variants in each consequence type category. To produce protein sequences, we focused on predicted consequences ‘missense variant’, ‘protein altering variant’, or ‘coding sequence variant’, and ignored variants on a transcript that were ‘non-coding transcript variant’ or ‘synonymous variant’. For other types of consequences we focused on those presenting more than 80 % pathogenic and likely pathogenic variants: ‘missense variant’, ‘protein altering variant’, ‘coding sequence variant’, ‘frameshift variant’, ‘splice donor variant’, ‘splice acceptor variant’, ‘stop retained’, ‘stop gained’, ‘inframe insertion’, and ‘inframe deletion’ (figure 2). The ‘stop lost’ consequences yielded different prevalence of pathogenic or likely pathogenic variants when considering the consequences reported by Ensembl vs. ClinVar (figure 2), which can be explained by the differences between Ensembl and ClinVar. Here, the Ensembl dataset consists of the exonic variants, but is not associated with any pathological phenotype. The ClinVar dataset, though, was not narrowed down to any particular regions in the genes of interest, but all the variants in it are associated with clinical conditions. Therefore, the Ensembl dataset is enriched with protein coding variants, whereas the ClinVar dataset is enriched with pathogenic variants. ‘stop lost’ consequences were included in our analysis. Altogether, the resulting table variants contained 69,256 unique variants located in 101 genes.

### Variant association with monogenic diabetes

We could map 2,701 of these variants to variants associated with monogenic diabetes according to ClinVar or the literature, termed *level 1* variants thereafter: 2,220 (82 %) mapped uniquely to ClinVar, 136 (5 %) to publications reviewed by Rafique et al., and 345 (13 %) to both (figure 4). An effect on protein sequences was predicted for 2,624 (97 %) of them, resulting in 12,643 different protein sequences when accounting for all isoforms. The reason for some variants not reaching the final translated sequences is that some transcripts in Ensembl do not have an associated canonical protein product. These are included when selecting exons in *Module 1* but do not yield protein sequences. On the other hand, genetic variation can cause translation of UTRs and other normally untranslated regions (22), and genetic variation in the UTRs and splice regions might affect the translation of the proteins in an indirect way (23,24), but the effects of these variants on amino acid sequences remain challenging to predict. We further filtered the variants to retain only the pathogenic and likely pathogenic variants, termed *level 2* variants thereafter, yielding 876 variants, of which 641 (73 %) mapped uniquely to ClinVar, 160 (18 %) to publications reviewed by Rafique et al., and 75 (9 %) to both (figure 4). An effect on protein sequences was predicted for 714 (82 %) of them, resulting in 3,776 distinct protein sequences.

**figure 4.**
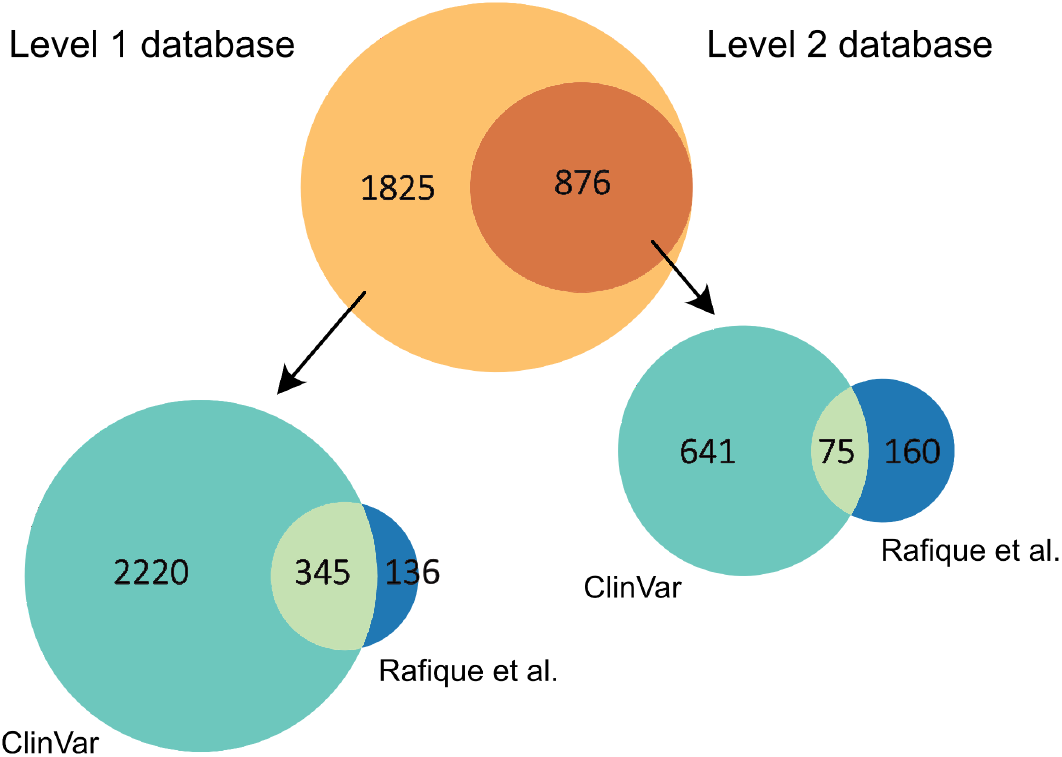
Venn diagram representing the number of variants in two levels of the database and the number of variants taken from different sources. *Level 1* database (at the left) consists of i. ClinVar variants associated with three phenotypes, i.e. “MODY”, “monogenic diabetes”, and “neonatal diabetes” mapped to Ensembl; ii. All variants from Rafique et al. mapped to Ensembl. *Level 2* database (at the right) consists of i. ClinVar “pathogenic” + “likely pathogenic” variants; ii. Variants from Rafique et al. excluded *BLK, KLF11*, and *PAX4*.

In both cases, the overlap between variants from ClinVar and the literature is limited. This can be explained by the fact that the ClinVar dataset consisted of the variants associated with all kinds of monogenic diabetes (101 genes) and the Rafique et al. dataset consisted only of the MODY variants (14 genes). Thus, we re-ran the pipeline considering only MODY variants from ClinVar. The overlap of the ClinVar MODY dataset was then 18 % and 10 % for the level 1 and 2 variants respectively. While the overlap was improved, it is still limited. This illustrates the importance to consider different sources of information when studying monogenic diabetes, and rare diseases in general.

It should be noted here that, in their work, Rafique et al. also used ClinVar as a source of variants. The variants from ClinVar that were not found by the literature text mining algorithm are listed in a separate supplementary file in their publication. We decided not to include this list in our work as we have included all the monogenic diabetes-associated variants from ClinVar anyway. This is another reason why the overlap of the variants taken from Rafique et al. and ClinVar seems limited in our analysis.

### Distribution of variants among the genes

We observed strong disparities in the number of variants associated with monogenic diabetes among the different genes. *HNF1A, GCK*, and *HNF1B* are the genes presenting most variants in both *level 1* and *level 2* databases, indicating that most of the monogenic diabetes-associated variants have been reported in these genes, as well as most of the ones confirmed as pathogenic or likely pathogenic (figure 3). Conversely, genes like *ABCC8* and *KCNJ11* feature many *level 1* variants, while only a few of those are confirmed to be pathogenic or likely pathogenic (figure 3). In fact, of all the genes observed, just about a third bear more than 20 variants, and only around 5 % have more than 100 variants. Figure 5 represents the number of gene variants in both levels of the database, where the number of variants in the gene is more than 20. The table 6 of the Supplementary materials and figure “number of variants per gene” in GitHub represent the number of variants in each gene in both levels of the database. Besides the biological reasons that some genes have more association with monogenic diabetes than others, this effect can be due to study biases or heterogeneous levels of information on these variants. For example, the most common genes known to cause MODY (*HNF1A, GCK, HNF1B*, and *HNF4A*)(2) might have received more attention than others (e.g. *ABCC8* and *KCNJ11*), yielding a lower confirmation rate for the pathogenicity of the variants. At the same time, *ABCC8* is reported to be also associated with other types of diabetes (25) and definitely deserves attention from the perspective of MODY associations as this may bring the researchers closer to the overall understanding of diabetes causal mechanisms and more precise treatment outcome predictions. The variant distribution in the studied genes can be observed in the figures visualizing the variants in the protein sequences (figure 3 and others in the “figure” directory at github.com/kuznetsovaks/MD_variants.git).

**figure 5.**
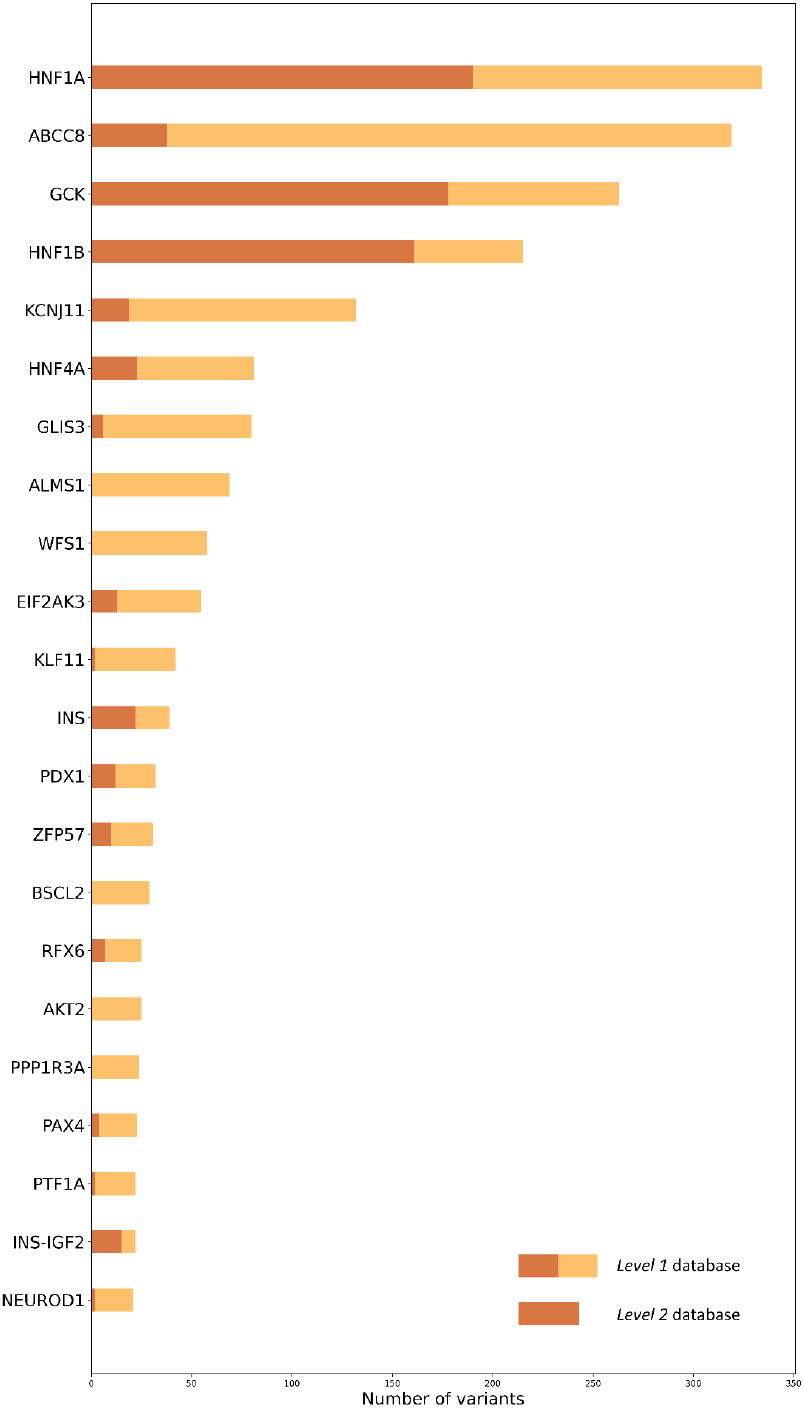
Number of variants in the genes with the largest number of variants. Selected are the genes in which the number of variants in the *level 1* database is more than 20. The full bars represent the number of variants in the *level 1* database, and the brown part of the bars represents the number of variants in the *level 2* database.

## Discussion

In this work, we presented a computational pipeline that allows for systematic monogenic diabetes-associated variant collection and mapping. The sources of information on the genetic variation are not unified which makes mapping of the variants challenging. An automated and reproducible pipeline for variant mapping has been developed and is available for public use. A database of variant protein sequences was created for the gene products of variants associated with monogenic diabetes. All known variants reported to be associated with monogenic diabetes published by the beginning of 2023 have been included in the database. The database contains variants with two levels of clinical significance: variants ever reported as associated with monogenic diabetes and pathogenic variants. Here we were considering the variants pathogenic if they had *pathogenic* or *likely pathogenic* clinical significance regardless of the star status according to ClinVar. All the monogenic diabetes-associated variants have been translated into protein products and can be compared to the canonical protein sequences. This will help predict the effect of genetic variation on the resulting protein structure and function.

The workflow is automated and aims to gather multiple variants from different sources and avoid their manual annotation. The implementation in Jupyter notebooks provides a good trade-off between automation and flexibility. For example, researchers can execute the entire pipeline as is, adapt it to specific use cases, execute only modules of interest, or completely change the set of genes to study another disease. The public availability, extensive documentation, and permissive license further enable the reuse of our work.

The collected genetic variants associated with monogenic diabetes have been translated to protein sequences and mapped to all known protein isoforms resulting in the collection of all predicted protein variant sequences. Our database is represented in the form of tables along with FASTA files and can be accessed both manually and automatically allowing implementation in various workflows. The localization of variants on proteins and protein domains can shed light on their possible consequences and pathogenicity. In turn, overlaying variant pathogenicity on protein sequences can help understanding protein function. The FASTA files can be supplemented with other proteins and used for proteomic search of mass spectrometry data. The variant protein sequences can also be used in protein structure modeling using tools like Alphafold (11).

This work illustrates how the different genes associated with monogenic diabetes show very different levels of annotation. Besides well-investigated genes, featuring a high number of variants with unambiguous consequences, many understudied genes bear variants lacking evidence of pathogenicity.

Furthermore, some variants simply lack basic genomic annotation, and are reported as amino acid changes, e.g. “Gly292Argfs”. An amino acid substitution cannot always be mapped to a single genetic variant. Furthermore, most of the genes encode several protein isoforms and knowing an amino acid change does not give information on which isoform it affects and how it maps to other isoforms. Thus, reporting single amino acid substitutions impairs their inclusion in genomic and bioinformatic studies. In this work, we manually curated these variants and were able to match some of them to genomic coordinates using other variants and aligning protein isoforms using the IsoAligner tool (26). For complex proteomic research in humans, it is important to account for common variation. In future work, we are going to combine our variant sequences with the database of human protein haplotypes (27).

Our work focused on single nucleotide variants (SNVs) or short indels and therefore does not cover larger insertion/deletion mutations causing monogenic diabetes. Large genomic rearrangements, such as full deletion of the *HNF1B* gene (28) or full deletion of 17q12 locus, have been shown to cause HNF1B-MODY (MODY5) and can be missed by conventional point mutation screening (28,29). These events cannot be directly translated to protein sequences and, as a result, are not reflected in our database.

In our work, we have analyzed the distribution of pathogenicity among the variants with known consequence types. Based on this analysis we have included variants classified as ‘splice donor variant’, ‘splice acceptor variant’ and filtered out the ‘splice region variants’. In future work, the question of splice region variant consequences should be given more attention as this is an understudied field, and these variants can play a significant role in rare diseases (30).

The research of monogenic diabetes is a dynamically developing field, and new variants are being constantly reported from different cohorts. Now with the pipeline we have developed, we and others can easily update MODY variant collections as new variants are reported. Our pipeline can further be used altogether or in parts to study other diseases. This can enable researchers to automatically and reproducibly collect variants associated with phenotypes of interest and consolidate them to a unified format. In research on rare diseases, the availability of flexible pipelines based on notebooks represents a good compromise between manual expert curation that lacks reproducibility and automated pipelines that cannot be tailored to the application.

## Supporting information

Supplementary file. Examples of the output.

## Abbreviations

API: application programming interface
MD: monogenic diabetes
MODY: maturity-onset diabetes of the young
ND: neonatal diabetes
OMIM: Online Mendelian Inheritance in Man
PMID: identifiers of scientific publications from the PubMed database
SNP: single nucleotide polymorphism
T2D: type 2 diabetes
UTR: untranslated region
VCF: Variant Calling Format
VEP: Variant Effect Predictor

## Acknowledgments

This work was supported by the Research Council of Norway (project #301178 to MV), the University of Bergen, and the Novo Nordisk Foundation (project NNF20OC0063872 to SJ).

This research was funded, in whole or in part, by the Research Council of Norway 301178. A CC BY or equivalent license is applied to any Author Accepted Manuscript (AAM) version arising from this submission, in accordance with the grant’s open access conditions.

## Data Availability

The source code of all the modules is freely available as Jupyter notebooks at github.com/kuznetsovaks/MD_variants.git and the resulting tables are deposited along with all the figures and the results in the same repository. Large FASTA files are available at doi.org/10.6084/m9.figshare.21444963.v2 All our code and data are openly available under a permissive CC-BY license.

## Contribution statement

