## Supplementary file. Examples of the output. for "A systematic mapping of the genomic and proteomic variation associated with monogenic diabetes"

### Supplementary materials

Ksenia Kuznetsova[1,$], Jakub Vašíček[1], Dafni Skiadopoulou[1], Janne Molnes[1,2], Miriam Udler[3], Stefan Johansson[1,2,], Pål Rasmus Njølstad[1,4], Alisa Manning[3,†], Marc Vaudel[1,5,6,†,$]

1. Mohn Center for Diabetes Precision Medicine, Department of Clinical Science, University of Bergen, Bergen, Norway
2. Department of Medical Genetics, Haukeland University Hospital, Bergen, Norway
3. Harvard Medical School, Mass General Hospital, Broad Institute, Cambridge and Boston, MA, USA
4. Children and Youth Clinic, Haukeland University Hospital, Bergen, Norway
5. Department of Genetics and Bioinformatics, Norwegian Institute of Public Health, Oslo, Norway
6. Computational Biology Unit (CBU), Department of Informatics, University of Bergen

[†] These authors jointly supervised the work
[$] To whom correspondence should be addressed

Table 1. Top rows of the reference Ensemble table (output of *Module 1*) shown as an example. This table is used as a reference to which all other tables are being mapped. The whole table can be downloaded from github.com/kuznetsovaks/MD_variants.git

| id | seq_region_name | start | end | strand | vf_allele | Location | coordinate | Gene | Transcript | Exon |
|---|---|---|---|---|---|---|---|---|---|---|
| rs1196762591 | 3 | 57227885 | 57227885 | 1 | A | 953801:05:00 | 3:57227885:T>A | ENSG00000157500 | ENST00000650354 | ENSE00003837686 |
| rs752732231 | 3 | 57227888 | 57227888 | 1 | A | 953801:08:00 | 3:57227888:C>A | ENSG00000157500 | ENST00000650354 | ENSE00003837686 |
| rs1559504574 | 3 | 57227889 | 57227892 | 1 | GGG | 953801:09:00 | 3:57227889:GGGG>GGG | ENSG00000157500 | ENST00000650354 | ENSE00003837686 |
| rs1022835366 | 3 | 57227890 | 57227890 | 1 | A | 953801:10:00 | 3:57227890:G>A | ENSG00000157500 | ENST00000650354 | ENSE00003837686 |
| rs1246196168 | 3 | 57227891 | 57227891 | 1 | A | 953801:11:00 | 3:57227891:G>A | ENSG00000157500 | ENST00000650354 | ENSE00003837686 |
| rs1180717064 | 3 | 57227896 | 57227896 | 1 | A | 953801:16:00 | 3:57227896:G>A | ENSG00000157500 | ENST00000650354 | ENSE00003837686 |
| rs1301549943 | 3 | 57227905 | 57227905 | 1 | T | 953801:25:00 | 3:57227905:C>T | ENSG00000157500 | ENST00000650354 | ENSE00003837686 |
| rs764047745 | 3 | 57227911 | 57227911 | 1 | T | 953801:31:00 | 3:57227911:G>T | ENSG00000157500 | ENST00000650354 | ENSE00003837686 |
| rs751342039 | 3 | 57227912 | 57227912 | 1 | T | 953801:32:00 | 3:57227912:A>T | ENSG00000157500 | ENST00000650354 | ENSE00003837686 |
| rs867398262 | 3 | 57227918 | 57227918 | 1 | A | 953801:38:00 | 3:57227918:C>A | ENSG00000157500 | ENST00000650354 | ENSE00003837686 |
| rs1171557373 | 3 | 57227930 | 57227930 | 1 | A | 953801:50:00 | 3:57227930:G>A | ENSG00000157500 | ENST00000650354 | ENSE00003837686 |
| rs1321589564 | 3 | 57227931 | 57227931 | 1 | A | 953801:51:00 | 3:57227931:C>A | ENSG00000157500 | ENST00000650354 | ENSE00003837686 |
| rs942686239 | 3 | 57227932 | 57227932 | 1 | T | 953801:52:00 | 3:57227932:C>T | ENSG00000157500 | ENST00000650354 | ENSE00003837686 |
| rs1329989662 | 3 | 57227933 | 57227933 | 1 | T | 953801:53:00 | 3:57227933:C>T | ENSG00000157500 | ENST00000650354 | ENSE00003837686 |
| rs1040198076 | 3 | 57227935 | 57227935 | 1 | T | 953801:55:00 | 3:57227935:C>T | ENSG00000157500 | ENST00000650354 | ENSE00003837686 |
| rs1317925966 | 3 | 57235582 | 57235582 | 1 | T | 953929:22:00 | 3:57235582:G>T | ENSG00000157500 | ENST00000650354 | ENSE00003577757 |
| rs770628528 | 3 | 57235584 | 57235584 | 1 | A | 953929:24:00 | 3:57235584:G>A | ENSG00000157500 | ENST00000650354 | ENSE00003577757 |
| rs756321385 | 3 | 57235591 | 57235591 | 1 | G | 953929:31:00 | 3:57235591:A>G | ENSG00000157500 | ENST00000650354 | ENSE00003577757 |
| rs1221003102 | 3 | 57235593 | 57235593 | 1 | T | 953929:33:00 | 3:57235593:G>T | ENSG00000157500 | ENST00000650354 | ENSE00003577757 |
| rs1377173261 | 3 | 57235596 | 57235596 | 1 | T | 953929:36:00 | 3:57235596:G>T | ENSG00000157500 | ENST00000650354 | ENSE00003577757 |
| rs368795074 | 3 | 57235599 | 57235599 | 1 | A | 953929:39:00 | 3:57235599:G>A | ENSG00000157500 | ENST00000650354 | ENSE00003577757 |
| rs753861823 | 3 | 57235602 | 57235602 | 1 | G | 953929:42:00 | 3:57235602:A>G | ENSG00000157500 | ENST00000650354 | ENSE00003577757 |
| rs772536998 | 3 | 57235604 | 57235604 | 1 | - | 953929:44:00 | 3:57235604:A>- | ENSG00000157500 | ENST00000650354 | ENSE00003577757 |

| | | | | | | | | | | |
|---|---|---|---|---|---|---|---|---|---|---|
| rs1343151977 | 3 | 57235608 | 57235608 | 1 | G | 953929:48:00 | 3:57235608:A>G | ENSG00000157500 | ENST00000650354 | ENSE00003577757 |
| rs776237801 | 3 | 57235615 | 57235615 | 1 | G | 953929:55:00 | 3:57235615:A>G | ENSG00000157500 | ENST00000650354 | ENSE00003577757 |
| rs779356223 | 3 | 57235617 | 57235617 | 1 | C | 953929:57:00 | 3:57235617:T>C | ENSG00000157500 | ENST00000650354 | ENSE00003577757 |
| rs188131823 | 3 | 57235618 | 57235618 | 1 | G | 953929:58:00 | 3:57235618:A>G | ENSG00000157500 | ENST00000650354 | ENSE00003577757 |
| rs1195723547 | 3 | 57235620 | 57235620 | 1 | G | 953930:00:00 | 3:57235620:A>G | ENSG00000157500 | ENST00000650354 | ENSE00003577757 |
| rs758848028 | 3 | 57235631 | 57235631 | 1 | A | 953930:11:00 | 3:57235631:G>A | ENSG00000157500 | ENST00000650354 | ENSE00003577757 |
| rs529656946 | 3 | 57235645 | 57235645 | 1 | G | 953930:25:00 | 3:57235645:A>G | ENSG00000157500 | ENST00000650354 | ENSE00003577757 |
| rs765656762 | 3 | 57235647 | 57235647 | 1 | T | 953930:27:00 | 3:57235647:C>T | ENSG00000157500 | ENST00000650354 | ENSE00003577757 |
| rs769557700 | 3 | 57235648 | 57235648 | 1 | A | 953930:28:00 | 3:57235648:G>A | ENSG00000157500 | ENST00000650354 | ENSE00003577757 |
| rs1232389093 | 3 | 57235654 | 57235654 | 1 | G | 953930:34:00 | 3:57235654:A>G | ENSG00000157500 | ENST00000650354 | ENSE00003577757 |
| rs775143822 | 3 | 57235656 | 57235656 | 1 | C | 953930:36:00 | 3:57235656:G>C | ENSG00000157500 | ENST00000650354 | ENSE00003577757 |
| rs748766381 | 3 | 57235659 | 57235659 | 1 | A | 953930:39:00 | 3:57235659:G>A | ENSG00000157500 | ENST00000650354 | ENSE00003577757 |
| rs1316373298 | 3 | 57237493 | 57237493 | 1 | G | 953961:13:00 | 3:57237493:A>G | ENSG00000157500 | ENST00000650354 | ENSE00003599120 |
| rs1310209188 | 3 | 57237502 | 57237502 | 1 | A | 953961:22:00 | 3:57237502:G>A | ENSG00000157500 | ENST00000650354 | ENSE00003599120 |
| rs1379418245 | 3 | 57237505 | 57237505 | 1 | T | 953961:25:00 | 3:57237505:C>T | ENSG00000157500 | ENST00000650354 | ENSE00003599120 |
| rs1241972240 | 3 | 57237511 | 57237511 | 1 | A | 953961:31:00 | 3:57237511:C>A | ENSG00000157500 | ENST00000650354 | ENSE00003599120 |

Table 2. Top rows of the ClinVar export for the phenotype “Monogenic diabetes” (input of Module 3) shown as an example. At this stage, the variants are not filtered and not mapped to Ensembl. The whole table along with the other two phenotypes used (i.e. “MODY” and “Neonatal diabetes”) can be downloaded from the “input” directory on github.com/kuznetsovaks/MD_variants.git

| Name | Gene(s) | Protein change | Condition(s) | Clinical significance (Last reviewed) | Review status | Accession | GRCh37 Chromosome | GRCh37 Location | GRCh38Chromosome | GRCh38Location | VariationID | AlleleID(s) | dbSNP ID | Canonical SPDI |
|---|---|---|---|---|---|---|---|---|---|---|---|---|---|---|
| NM_000352.6(ABCC8):c.1127C>T (p.Ser376Phe) | ABCC8 | S375F, S376F | Monogenic diabetes\|Transitory neonatal diabetes mellitus\|Maturity onset diabetes mellitus in young | Uncertain significance(Last reviewed: Jul 6, 2018) | criteria provided, multiple submitters, no conflicts | VCV000917422 | 11 | 17474715 | 11 | 17453168 | 917422 | 905790 | rs777461294 | NC_000011.10:17453167:G:A |
| NM_000352.6(ABCC8):c.1874C>T (p.Ala625Val) | ABCC8 | A625V, A624V | not provided\|not specified\|Monogenic diabetes | Conflicting interpretations of pathogenicity(Last reviewed: Apr 2, 2018) | criteria provided, conflicting interpretations | VCV000596784 | 11 | 17450161 | 11 | 17428614 | 596784 | 587845 | rs148709148 | NC_000011.10:17428613:G:A |
| NM_000352.6(ABCC8):c.4136G>A (p.Arg1379His) | ABCC8 | R1379H, R1380H, R1378H, R1401H | Monogenic diabetes\|not provided | Likely pathogenic(Last reviewed: Jan 17, 2022) | criteria provided, multiple submitters, no conflicts | VCV000585346 | 11 | 17417461 | 11 | 17395914 | 585346 | 577178 | rs193922401 | NC_000011.10:17395913:C:T |
| NM_000352.6(ABCC8):c.2666A>C (p.Lys889Thr) | ABCC8 | K889T, K890T, K911T, K888T | Monogenic diabetes\|not specified\|Diabetes mellitus, transient neonatal, 2\|Hyperinsulinemic hypoglycemia, familial, 1\|Permanent neonatal diabetes mellitus | Uncertain significance(Last reviewed: Dec 29, 2021) | criteria provided, multiple submitters, no conflicts | VCV000558717 | 11 | 17432091 | 11 | 17410544 | 558717 | 546501 | rs761862121 | NC_000011.10:17410543:T:G |
| NM_000352.6(ABCC8):c.250G>A (p.Val84Ile) | ABCC8 | V84I | Monogenic diabetes\|Hyperinsulinemic hypoglycemia, familial, 1 | Conflicting interpretations of pathogenicity(Last reviewed: Mar 9, 2022) | criteria provided, conflicting interpretations | VCV000554707 | 11 | 17496473 | 11 | 17474926 | 554707 | 546216 | rs775776658 | NC_000011.10:17474925:C:T |

| | | | | | | | | | | | | | | |
|---|---|---|---|---|---|---|---|---|---|---|---|---|---|---|
| NM_000352.6(ABCC8):c.3817A>T (p.Arg1273Trp) | ABCC8 | R1273W, R1274W, R1295W, R1272W | Transitory neonatal diabetes mellitus\|Monogenic diabetes\|Maturity onset diabetes mellitus in young | Uncertain significance(Last reviewed: Aug 1, 2017) | criteria provided, multiple submitters, no conflicts | VCV000549540 | 11 | 17419281 | 11 | 17397734 | 549540 | 540023 | rs1554906389 | NC_000011.10:17397733:T:A |
| NM_000352.6(ABCC8):c.4043A>G (p.Asn1348Ser) | ABCC8 | N1348S, N1349S, N1370S, N1347S | Transitory neonatal diabetes mellitus\|Monogenic diabetes\|Maturity onset diabetes mellitus in young | Uncertain significance(Last reviewed: May 19, 2017) | criteria provided, multiple submitters, no conflicts | VCV000549539 | 11 | 17418539 | 11 | 17396992 | 549539 | 540022 | rs1554905775 | NC_000011.10:17396991:T:C |
| NM_000352.6(ABCC8):c.2992C>T (p.Arg998Ter) | ABCC8 | R998*, R999*, R1020*, R997* | Hereditary hyperinsulinism\|not provided\|Monogenic diabetes\|Hyperinsulinemic hypoglycemia, familial, 1 | Conflicting interpretations of pathogenicity(Last reviewed: Dec 9, 2021) | criteria provided, conflicting interpretations | VCV000434053 | 11 | 17428605 | 11 | 17407058 | 434053 | 429204 | rs769518471 | NC_000011.10:17407057:G:A |
| NM_000352.6(ABCC8):c.4563G>T (p.Lys1521Asn) | ABCC8 | K1521N, K1522N, K1520N, K1543N | Monogenic diabetes\|Transitory neonatal diabetes mellitus\|not specified\|Maturity onset diabetes mellitus in young\|not provided | Conflicting interpretations of pathogenicity(Last reviewed: Apr 26, 2021) | criteria provided, conflicting interpretations | VCV000434043 | 11 | 17415289 | 11 | 17393742 | 434043 | 429195 | rs142272833 | NC_000011.10:17393741:C:A |
| NM_000352.6(ABCC8):c.375C>G (p.His125Gln) | ABCC8 | H125Q | not provided\|not specified\|Monogenic diabetes\|Hyperinsulinemic hypoglycemia, familial, 1\|Diabetes mellitus, transient neonatal, 2\|Permanent neonatal diabetes mellitus | Conflicting interpretations of pathogenicity(Last reviewed: May 11, 2022) | criteria provided, conflicting interpretations | VCV000393430 | 11 | 17491685 | 11 | 17470138 | 393430 | 380249 | rs60637558 | NC_000011.10:17470137:G:C |
| NM_000352.6(ABCC8):c.1067A>G (p.Tyr356Cys) | ABCC8 | Y356C, Y355C | not specified\|Hereditary hyperinsulinism\|Monogenic diabetes\|not provided\|Permanent neonatal diabetes | Conflicting interpretations of pathogenicity(Last reviewed: May 3, 2022) | criteria provided, conflicting interpretations | VCV000393429 | 11 | 17474775 | 11 | 17453228 | 393429 | 380248 | rs59852838 | NC_000011.10:17453227:T:C |

| | | | | | | | | | | | | | |
|---|---|---|---|---|---|---|---|---|---|---|---|---|---|
| | | | mellitus\|Diabetes mellitus, transient neonatal, 2\|Hyperinsulinemic hypoglycemia, familial, 1 | | | | | | | | | | |
| NM_000352.6(ABCC8):c.3875A>G (p.Asn1292Ser) | ABCC8 | N1292S, N1293S, N1291S, N1314S | Monogenic diabetes\|Transitory neonatal diabetes mellitus\|Maturity onset diabetes mellitus in young | Uncertain significance(Last reviewed: Aug 16, 2016) | criteria provided, multiple submitters, no conflicts | VCV000393428 | 11 | 17418853 | 11 | 17397306 | 393428 | 380247 | rs763104338 | NC_000011.10:17397305:T:C |
| NM_000352.6(ABCC8):c.3976G>A (p.Glu1326Lys) | ABCC8 | E1326K, E1327K, E1348K, E1325K | Monogenic diabetes\|Hyperinsulinemic hypoglycemia, familial, 1\|not provided | Uncertain significance(Last reviewed: Nov 4, 2021) | criteria provided, multiple submitters, no conflicts | VCV000288731 | 11 | 17418752 | 11 | 17397205 | 288731 | 272968 | rs200563930 | NC_000011.10:17397204:C:T |
| NM_000352.6(ABCC8):c.1858C>T (p.Arg620Cys) | ABCC8 | R620C, R619C | not provided\|Monogenic diabetes\|Hereditary hyperinsulinism\|not specified\|Hyperinsulinemic hypoglycemia, familial, 1 | Conflicting interpretations of pathogenicity(Last reviewed: May 3, 2021) | criteria provided, conflicting interpretations | VCV000210071 | 11 | 17450177 | 11 | 17428630 | 210071 | 207849 | rs58241708 | NC_000011.10:17428629:G:A |
| NM_000352.6(ABCC8):c.886G>A (p.Gly296Arg) | ABCC8 | G296R, G295R | Hereditary hyperinsulinism\|Monogenic diabetes\|not specified\|Hyperinsulinemic hypoglycemia, familial, 1 | Uncertain significance(Last reviewed: Nov 16, 2021) | criteria provided, multiple submitters, no conflicts | VCV000035624 | 11 | 17482160 | 11 | 17460613 | 35624 | 44289 | rs148529020 | NC_000011.10:17460612:C:T |
| NM_000352.6(ABCC8):c.4135C>A (p.Arg1379Ser) | ABCC8 | R1379S, R1380S, R1401S, R1378S | Hereditary hyperinsulinism\|Monogenic diabetes\|not provided\|not specified\|Hyperinsulinemic hypoglycemia, familial, 1 | Uncertain significance(Last reviewed: Jan 22, 2020) | criteria provided, multiple submitters, no conflicts | VCV000035614 | 11 | 17417462 | 11 | 17395915 | 35614 | 44279 | rs137852673 | NC_000011.10:17395914:G:T |

Table 3. Top rows of the ClinVar mapped table containing indels (output of *Module 3*). This table is further used for VCF. The whole table can be downloaded from github.com/kuznetsovaks/MD_variants.git

| chrom | pos | ref | alt | accession |
|---|---|---|---|---|
| 12 | 121001081 | GG | G | VCV001700003 |
| 12 | 121001136 | AA | | VCV001687087 |
| 12 | 121001116 | AGAG | AG | VCV001687086 |
| 12 | 121001098 | C | | VCV001687084 |
| 12 | 121001158 | CATCTCCACCCAGATGGCCTCTTCCTCC | CATCTCCACCCAGATGGCCTCTTCCTCCATCTCCACCCAGATGGCCTCTTCCTCC | VCV001687077 |
| 12 | 121001068 | CC | | VCV001687070 |
| 7 | 44145499 | GGGG | GGG | VCV001526254 |
| 7 | 44145579 | TGATGA | TGA | VCV001303094 |
| 7 | 44145255 | CGC | C | VCV001301411 |
| 7 | 44153361 | GGG | GG | VCV001254640 |
| 7 | 44149980 | GATAG | GATAGATAG | VCV001195505 |
| 12 | 120988910 | A | | VCV001700001 |
| 12 | 120988883 | ACAACA | ACA | VCV001693218 |
| 12 | 120999577 | CC | C | VCV001693217 |
| 12 | 120994316 | CC | T | VCV001691371 |
| 12 | 120978897 | CC | A | VCV001687081 |
| 12 | 120978906 | GGGGGAGTCCTGCGGCGG | G | VCV001677040 |
| 12 | 120994311 | GGG | GGGG | VCV001676719 |
| 12 | 120994260 | CC | C | VCV001676715 |
| 12 | 120978483 | TC | | VCV001676714 |
| 12 | 120978971 | GGGG | GGG | VCV001676691 |
| 12 | 120978959 | GGGGG | GGGGGG | VCV001676689 |

Table 4. Top rows of the table containing SNVs from ClinVar mapped to the Ensemble reference table (output of *Module 3*). This table is further used for VCF. The whole table can be downloaded from github.com/kuznetsovaks/MD_variants.git. This kind of table is also produced by *Modules 4* and *5* and altogether passed to *Module 6*.

| chrom | pos | ref | alt | accession |
|---|---|---|---|---|
| 3 | 57238111 | G | A | rs796065047 |
| 3 | 57260016 | T | A | rs869320673 |
| 7 | 113918369 | G | A | rs141223649 |
| 7 | 113918581 | G | C | rs1057524893 |
| 7 | 113918700 | G | A | rs115949425 |
| 7 | 113918864 | C | T | rs8192687 |
| 7 | 113877832 | A | C | rs139484221 |
| 7 | 113877833 | T | C | rs77456357 |
| 7 | 113878048 | T | C | rs149701175 |
| 7 | 113878102 | A | C | rs35572169 |
| 7 | 113878120 | C | T | rs374950521 |
| 7 | 113878207 | C | T | rs148408134 |

Table 5. Top rows of the table containing variants from the final variant database mapped to the transcript sequences from Ensemble (one of the outputs of Module “Translation”). This mapping is dome by the tools from the ProVar package (github.com/ProGenNo/ProHap): Tho whole table is available in the “results” directory at github.com/kuznetsovaks/MD_variants.gitt

| transcriptID | chromosome | transcript_biotype | variantID | DNA_change | cDNA_change | protein_change | reading_frame | protein_prefix_length | start_missing | start_lost | splice_site_affected |
|---|---|---|---|---|---|---|---|---|---|---|---|
| ENST00000493373 | 1 | protein_coding | ENST00000493373_MDvar_rs71653619_G>A | 7970934:G>A | 348:G>A | 97:R>97:Q | 2 | 18 | TRUE | FALSE | |
| ENST00000493678 | 1 | protein_coding | ENST00000493678_MDvar_rs71653619_G>A | 7970934:G>A | 359:G>A | 97:R>97:Q | 1 | 22 | TRUE | FALSE | |
| ENST00000377493 | 1 | protein_coding | ENST00000377493_MDvar_rs71653619_G>A | 7970934:G>A | 290:G>A | 77:R>77:Q | 1 | 19 | TRUE | FALSE | |
| ENST00000338639 | 1 | protein_coding | ENST00000338639_MDvar_rs71653619_G>A | 7970934:G>A | 398:G>A | 97:R>97:Q | 1 | 35 | TRUE | FALSE | |
| ENST00000377491 | 1 | protein_coding | ENST00000377491_MDvar_rs71653619_G>A | 7970934:G>A | 503:G>A | 97:R>97:Q | 1 | 70 | TRUE | FALSE | |
| ENST00000377488 | 1 | protein_coding | ENST00000377488_MDvar_rs71653619_G>A | 7970934:G>A | 448:G>A | 97:R>97:Q | 0 | 52 | TRUE | FALSE | |
| ENST00000371059 | 1 | protein_coding | ENST00000371059_MDvar_rs35573508_A>T | 65572326:A>T | 555:A>T | 123:D>123:V | 2 | 61 | TRUE | FALSE | 4 |
| ENST00000371059 | 1 | protein_coding | ENST00000371059_MDvar_rs35573508_A>G | 65572326:A>G | 555:A>G | 123:D>123:G | 2 | 61 | TRUE | FALSE | 4 |
| ENST00000371059 | 1 | protein_coding | ENST00000371059_MDvar_rs147667928_G>T | 65572385:G>T | 614:G>T | 143:V>143:L | 2 | 61 | TRUE | FALSE | |
| ENST00000371059 | 1 | protein_coding | ENST00000371059_MDvar_rs148349369_G>A | 65592820:G>A | 842:G>A | 219:V>219:I | 2 | 61 | TRUE | FALSE | |
| ENST00000371059 | 1 | protein_coding | ENST00000371059_MDvar_rs1057524881_G>T | 65601452:G>T | 1239:G>T | 351:C>351:F | 2 | 61 | TRUE | FALSE | |
| ENST00000371059 | 1 | protein_coding | ENST00000371059_MDvar_rs751025808_A>G | 65601536:A>G | 1323:A>G | 379:Q>379:R | 2 | 61 | TRUE | FALSE | |
| ENST00000371059 | 1 | protein_coding | ENST00000371059_MDvar_rs937591370_A>G | 65601849:A>G | 1476:A>G | 430:N>430:S | 2 | 61 | TRUE | FALSE | |

Table 6. The numbers of variants in each gene in the *level 1* and *level 2* databases.

| gene_name | number of variants level 1 | number of variants level 2 | gene_name | number of variants level 1 | number of variants level 2 | gene_name | number of variants level 1 | number of variants level 2 | gene_name | number of variants level 1 | number of variants level 2 |
|---|---|---|---|---|---|---|---|---|---|---|---|
| *HNF1A* | 334 | 190 | *PTF1A* | 22 | 2 | *APPL1* | 3 | 2 | *PAX2* | 1 | 0 |
| *ABCC8* | 319 | 38 | *INS-IGF2* | 22 | 15 | *GLUD1* | 3 | 0 | *ITGB3* | 1 | 0 |
| *GCK* | 263 | 178 | *NEUROD1* | 21 | 2 | *FBN1* | 3 | 2 | *FOXP1* | 1 | 0 |
| *HNF1B* | 215 | 161 | *C12orf43* | 20 | 7 | *PPARG* | 3 | 0 | *GATA4* | 1 | 0 |
| *KCNJ11* | 132 | 19 | *CEL* | 15 | 3 | *MECP2* | 2 | 0 | *EDEM2* | 1 | 0 |
| *HNF4A* | 81 | 23 | *AGPAT2* | 13 | 0 | *EFL1* | 2 | 0 | *GPR161* | 1 | 1 |
| *GLIS3* | 80 | 6 | *LEPR* | 13 | 0 | *LEP* | 2 | 1 | *GRIN2B* | 1 | 0 |
| *ALMS1* | 69 | 0 | *INSR* | 12 | 0 | *LAMA2* | 2 | 0 | *HBB* | 1 | 0 |
| *WFS1* | 58 | 0 | *BLK* | 12 | 3 | *PURA* | 2 | 1 | *IER3IP1* | 1 | 0 |
| *EIF2AK3* | 55 | 13 | *PLIN1* | 11 | 0 | *PAX6* | 2 | 2 | *KCNH2* | 1 | 0 |
| *KLF11* | 42 | 2 | *MC4R* | 9 | 0 | *CAV1* | 2 | 0 | *PARK7* | 1 | 0 |
| *INS* | 39 | 22 | *HADH* | 9 | 0 | *SHLD2* | 2 | 0 | *KCNQ1* | 1 | 1 |
| *PDX1* | 32 | 12 | *GATA6* | 8 | 0 | *PRKAG2* | 1 | 0 | *ASB14* | 1 | 0 |
| *ZFP57* | 31 | 10 | *SLC2A2* | 8 | 1 | *SCN1A* | 1 | 1 | *KMT2E* | 1 | 0 |
| *BSCL2* | 29 | 0 | *SIM1* | 7 | 0 | *TRIP11* | 1 | 1 | *MAGEL2* | 1 | 1 |
| *RFX6* | 25 | 7 | *FOXP3* | 6 | 1 | *SHANK3* | 1 | 1 | *MBL2* | 1 | 0 |
| *AKT2* | 25 | 0 | *LMNA* | 6 | 0 | *PDIA6* | 1 | 0 | *MLKL* | 1 | 1 |
| *PPP1R3A* | 24 | 0 | *CAVIN1* | 6 | 0 | *RET* | 1 | 0 | *NLRP3* | 1 | 0 |
| *PAX4* | 23 | 4 | *SLC19A2* | 6 | 0 | *LZTR1* | 1 | 1 | *KCNQ2* | 1 | 1 |

FASTA 1. Example of FASTA entries each containing the protein description and the variant protein sequence (output of the module “Translation”). This FASTA is producedby  the tools from the ProVar package (github.com/ProGenNo/ProHap): Tho whole FASTA file is available at doi.org/10.6084/m9.figshare.21444963.v1

>generic_var|prot_0|position_within_protein:10 protein_IDs:MDvar_d38 start:10 matching_proteins:ENST00000644772_MDvar_rs547150342_C>T reading_frame:1
MPLAFCGSENHSAAYRVDQGVLNNGCFVDALNVVPHVFLLFITFPILFIGWGSQSSKVHIHHSTWLHFPGHNLRWILTFMLLFVLVCEIAEGILSDGVTE
SHHLHLYMPAGMAFMAAVTSVVYYHNIETSNFPKLLIALLVYWTLAFITKTIKFVKFLDHAIGFSQLRFCLTGLLVILYGMLLLVEVNVIRVRRYIFFKTPRE
VKPPEDLQDLGVRFLQPFVNLLSKGTYWWMNAFIKTAHKKPIDLRAIGKLPIAMRALTNYQRLCEAFDAQVRKDIQGTQGARAIWQALSHAFGRRLVL
SSTFRILADLLGFAGPLCIFGIVDHLGKENDVFQPKTQFLGVYFVSSQEFLANAYVLAVLLFLALLLQRTFLQASYYVAIETGINLRGAIQTKIYNKIMHLST
SNLSMGEMTAGQICNLVAIDTNQLMWFFFLCPNLWAMPVQIIVGVILLYYILGVSALIGAAVIILLAPVQYFVATKLSQAQRSTLEYSNERLKQTNEMLRGI
KLLKLYAWENIFRTRVETTRRKEMTSLRAFAIYTSISIFMNTAIPIAAVLITFVGHVSFFKEADFSPSVAFASLSLFHILVTPLFLLSSVVRSTVKALVSVQKL
SEFLSSAEIREEQCAPHEPTPQGPASKYQAVEEGLPSLHSSQPSGSSPPNHPQPLRVVNRKRPAREDCRGLTGPLQSLVPSADGDADNCCVQIMGG
YFTWTPDGIPTLSNITIRIPRGQLTMIVGQVGCGKSSLLLAALGEMQKVSGAVFWSSLPDSEIGEDPSPERETATDLDIRKRGPVAYASQKPWLLNATVE
ENIIFESPFNKQRYKMVIEACSLQPDIDILPHGDQTQIGERGINLSGGQRQRISVAQALYQHANVVFLDDPFSALDIHLSDHLMQAGILELLRDDKRTVVL
VTHKLQYLPHADWIIAMKDGTIQREGTLKDFQRSECQLFEHWKTLMNRQDQELEKETVTERKATEPPQGLSRAMSSRDGLLQDEEEEEEEAAESEE
DDNLSSMLHQRAEIPWRACAKYLSSAGILLLSLLVFSQLLKHMVLVAIDYWLAKWTDSALTLTPAARNCSLSQECTLDQTVYAMVFTVLCSLGIVLCLVT
SVTVEWTGLKVAKRLHRSLLNRIILAPMRFFETTPLGSILNRFSSDCNTIDQHIPSTLECLSRSTLLCVSALAVISYVTPVFLVALLPLAIVCYFIQKYFRVA
SRDLQQLDDTTQLPLLSHFAETVEGLTTIRAFRYEARFQQKLLEYTDSNNIASLFLTAANRWLEVRMEYIGACVVLIAAVTSISNSLHRELSAGLVGLGLT
YALMVSNYLNWMVRNLADMELQLGAVKRIHGLLKTEAESYEGLLAPSLIPKNWPDQGKIQIQNLSVRYDSSLKPVLKHVNALIAPGQKIGICGRTGSGK
SSFSLAFFRMVDTFEGHIIIDGIDIAKLPLHTLRSRLSIILQDPVLFSGTIRFNLDPERKCSDSTLWEALEIAQLKLVVKALPGGLDAIITEGGENFSQGQRQ
LFCLARAFVRKTSIFIMDEATASIDMATENILQKVVMTAFADRTVVTIAHRVHTILSADLVIVLKRGAILEFDKPEKLLSRKDSVFASFVRADK

>generic_var|prot_0_REVERSED|position_within_protein:10 protein_IDs:MDvar_d38 start:10
matching_proteins:ENST00000644772_MDvar_rs547150342_C>T reading_frame:1_REVERSED
KDARVFSAFVSDKRSLLKEPKDFELIAGRKLVIVLDASLITHVRHAITVVTRDAFATMVVKQLINETAMDISATAEDMIFISTKRVFARALCFLQRQGQSFN
EGGETIIADLGGPLAKVVLKLQAIELAEWLTSDSCKREPDLNFRITGSFLVPDQLIISLRSRLTHLPLKAIDIGDIIIHGEFTDVMRFFALSFSSKGSGTRGCI
GIKQGPAILANVHKLVPKLSSDYRVSLNQIQIKGQDPWNKPILSPALLGEYSEAETKLLGHIRKVAGLQLEMDALNRVMWNLYNSVMLAYTLGLGVLGA
SLERHLSNSISTVAAILVVCAGIYEMRVELWRNAATLFLSAINNSDTYELLKQQFRAEYRFARITTLGEVTEAFHSLLPLQTTDDLQQLDRSAVRFYKQIF
YCVIALPLLAVLFVPTVYSIVALASVCLLTSRSLCELTSPIHQDITNCDSSFRNLISGLPTTEFFRMPALIIRNLLSRHLRKAVKLGTWEVTVSTVLCLVIGLS

CLVTFVMAYVTQDLTCEQSLSCNRAAPTLTLASDTWKALWYDIAVLVMHKLLQSFVLLSLLLIGASSLYKACARWPIEARQHLMSSLNDDEESEAAEEE
EEEEDQLLGDRSSMARSLGQPPETAKRETVTEKELEQDQRNMLTKWHEFLQCESRQFDKLTGERQITGDKMAIIWDAHPLYQLKHTVLVVTRKDDRL
LELIGAQMLHDSLHIDLASFPDDLFVVNAHQYLAQAVSIRQRQGGSLNIGREGIQTQDGHPLIDIDPQLSCAEIVMKYRQKNFPSEFIINEEVTANLLWPK
QSAYAVPGRKRIDLDTATEREPSPDEGIESDPLSSWFVAGSVKQMEGLAALLLSSKGCGVQGVIMTLQGRPIRITINSLTPIGDPTWTFYGGMIQVCCN
DADGDASPVLSQLPGTLGRCDERAPRKRNVVRLPQPHNPPSSGSPQSSHLSPLGEEVAQYKSAPGQPTPEHPACQEERIEASSLFESLKQVSVLAK
VTSRVVSSLLFLPTVLIHFLSLSAFAVSPSFDAEKFFSVHGVFTILVAAIPIATNMFISISTYIAFARLSTMEKRRTTEVRTRFINEWAYLKLLKIGRLMENTQ
KLRENSYELTSRQAQSLKTAVFYQVPALLIIVAAGILASVGLIYYLLIVGVIIQVPMAWLNPCLFFFWMLQNTDIAVLNCIQGATMEGMSLNSTSLHMIKNYI
KTQIAGRLNIGTEIAVYYSAQLFTRQLLLALFLLVALVYANALFEQSSVFYVGLFQTKPQFVDNEKGLHDVIGFICLPGAFGLLDALIRFTSSLVLRRGFAH
SLAQWIARAGQTGQIDKRVQADFAECLRQYNTLARMAIPLKGIARLDIPKKHATKIFANMWWYTGKSLLNVFPQLFRVGLDQLDEPPKVERPTKFFIYR
RVRIVNVEVLLLMGYLIVLLGTLCFRLQSFGIAHDLFKVFKITKTIFALTWYVLLAILLKPFNSTEINHYYVVSTVAAMFAMGAPMYLHLHHSETVGDSLIG
EAIECVLVFLLMFTLIWRLNHGPFHLWTSHHIHVKSSQSGWGIFLIPFTIFLLFVHPVVNLADVFCGNNLVGQDVRYAASHNESGCFALPM

>generic_ensref|prot_2|position_within_protein:0 protein_IDs:ENSP00000255224.2 start:0 matching_proteins:ENSP00000255224.2
reading_frame:-
MAPITTSREEFDEIPTVVGIFSAFGLVFTVSLFAWICCQRKSSKSNKTPPYKFVHVLKGVDIYPENLNSKKKFGADDKNEVKNKPAVPKNSLHLDLEKRD
LNGNFPKTNLKPGSPSDLENATPKLFLEGEKESVSPESLKSSTSLTSEEKQEKLGTLFFSLEYNFERKAFVVNIKEARGLPAMDEQSMTSDPYIKMTIL
PEKKHKVKTRVLRKTLDPAFDETFTFYGIPYTQIQELALHFTILSFDRFSRDDIIGEVLIPLSGIELSEGKMLMNREIIKRNVRKSSGRGELLISLCYQSTTN
TLTVVVLKARHLPKSDVSGLSDPYVKVNLYHAKKRISKKKTHVKKCTPNAVFNELFVFDIPCEGLEDISVEFLVLDSERGSRNEVIGQLVLGAAAEGTGG
EHWKEICDYPRRQIAKWHVLCDG

>generic_ensref|prot_2_REVERSED|position_within_protein:0 protein_IDs:ENSP00000255224.2 start:0 matching_proteins:ENSP00000255224.2
reading_frame:-_REVERSED
GDCLVHWKAIQRRPYDCIEKWHEGGTGEAAAGLVLQGIVENRSGRESDLVLFEVSIDELGECPIDFVFLENFVANPTCKKVHTKKKSIRKKAHYLNVKV
YPDSLGSVDSKPLHRAKLVVVTLTNTTSQYCLSILLEGRGSSKRVNRKIIERNMLMKGESLEIGSLPILVEGIIDDRSFRDFSLITFHLALEQIQTYPIGYFT
FTEDFAPDLTKRLVRTKVKHKKEPLITMKIYPDSTMSQEDMAPLGRAEKINVVFAKREFNYELSFFLTGLKEQKEESTLSTSSKLSEPSVSEKEGELFLK
PTANELDSPSGPKLNTKPFNGNLDRKELDLHLSNKPVAPKNKVENKDDAGFKKKSNLNEPYIDVGKLVHVFKYPPTKNSKSSKRQCCIWAFLSVTFVL
GFASFIGVVTPIEDFEERSTTIPAM
